# Dual signaling of Wamide myoinhibitory peptides through a peptide-gated channel and a GPCR in *Platynereis*

**DOI:** 10.1101/261990

**Authors:** Axel Schmidt, Philipp Bauknecht, Elizabeth A. Williams, Katrin Augustinowski, Stefan Gründer, Gáspár Jékely

## Abstract

Neuropeptides commonly signal by metabotropic G-protein coupled receptors (GPCRs). In some mollusks and cnidarians, RFamide neuropeptides mediate fast ionotropic signaling by peptide-gated ion channels that belong to the DEG/ENaC family. Here we describe a neuropeptide system with a dual mode of signaling by both a peptide-gated channel and a GPCR. We identified and characterised a peptide-gated channel in the marine annelid *Platynereis dumerilii* that is specifically activated by Wamide myoinhibitory peptides derived from the same proneuropeptide. The myoinhibitory peptide-gated ion channel (MGIC) belongs to the DEG/ENaC family and is paralogous to RFamide-gated channels. *Platynereis* myoinhibitory peptides also activate a previously described GPCR, MAG. We measured the potency of all Wamides on both MGIC and MAG and identified peptides that preferentially activate one or the other receptor. Analysis of a single-cell transcriptome resource indicates that MGIC and MAG signal to distinct target neurons. The identification of a Wamide-gated channel suggests that peptide-gated channels are more diverse and widespread in animals than previously appreciated. The possibility of neuropeptide signaling by both ionotropic and metabotropic receptors to different target cells in the same organism highlights an additional level of complexity in peptidergic signaling networks.

## Introduction

Neuropeptides represent a large and diverse class of neuronal signaling molecules with a wide range of functions in the regulation of behavior, physiology, and development. Neuropeptides can be released at synapses (either alone or together with small molecule transmitters (1) and act on postsynaptic neurons. They can also be released extra-synaptically and act via volume transmission in the nervous system (2). Release at neurohaemal release sites allows neuropeptides to act as hormones in long-range signaling. The vast majority of neuropeptides signal through G-protein coupled receptors (GPCRs) and alter target-cell physiology by metabotropic signals mediated by G-proteins (3).

In some mollusks and cnidarians, RFamide neuropeptides can mediate fast ionotropic signaling through peptide-gated ion channels. The first physiological evidence that an ion channel can be directly activated by a neuropeptide (FMRFamide) was obtained in the snail *Helix* (4). This FMRFamide-activated Na^+^ channel (FaNaCh) could be blocked by amiloride and was shown to belong to the DEG/ENaC family of ion channels (5, 6). Ion channels directly gated by RFamide neuropeptides were later discovered in *Hydra magnipapillata*, a fresh-water cnidarian (7–11). The *Hydra* sodium channels (HyNaC) also belong to the DEG/ENaC gene family but are only distantly related to mollusk FaNaCs. Their discovery suggested that RFamide neuropeptides may have ancestrally signaled by an ion channel to mediate fast neuronal signaling. However, the sporadic phylogenetic distribution of the characterized peptide-gated channels suggested that peptide-gated ionotropic receptors have limited significance in neuronal communication across animal diversity and may be restricted to RFamide-gated channels in a few animal groups.

To further explore the phylogenetic distribution and diversity of peptide-gated channels, we set out to identify such channels in the marine annelid *Platynereis dumerilii. Platynereis* has emerged as a useful model in recent years for the study of neuroendocrine signaling and neuropeptide evolution. A comprehensive complement of neuropeptides has been described for this species, consisting of 100 proneuropeptide genes (12). Several of these neuropeptides have been mapped to the nervous system and have been studied functionally (13–17). Furthermore, over 20 different neuropeptide and monoamine GPCRs have been experimentally identified in *Platynereis*, including the founding members of several major bilaterian neuropeptide families (achatin, FMRFamide, RGWamide, FLamide, and elevenin) and the first invertebrate adrenergic receptors (17–19).

Here we identified an ion channel that is gated by *Platynereis* myoinhibitory peptides (MIPs). Previously, a GPCR specifically activated by *Platynereis* MIPs has been identified (15). The identification of a MIP-gated channel provides an example where neuropeptides from the same precursor can signal both by an ionotropic and a metabotropic receptor. Our findings indicate that peptide-gated channels are more widespread and diverse than previously thought.

## Results

### *Platynereis* DEG/ENaC channels define a family paralogous to FaNaCs

Based on sequence homology to the *Helix aspersa* FaNaC receptor we cloned four DEG/EnaC ion channel subunits of the annelid worm *Platynereis dumerilii*. The *Platynereis* DEG/ENaCs (referred to as “ENaCs”) were between 508-614 amino acids long (ENaC4: 614, ENaC5: 589, ENaC6: 591, and ENaC7: 508) and shared a sequence identity of 32-59% among each other.

To analyze the phylogenetic position of *Platynereis* ENaCs, we performed clustering and phylogenetic analyses. Sequence-similarity-based clustering revealed that *Platynereis* ENaC4-7 form a cluster with other lophotrochozoan sequences, including the mollusk FaNaCs (Figure 1A). This cluster is related to vertebrate ENaCs and is more distantly related to acid-sensing ion channels (ASICs). A maximum likelihood phylogenetic analysis of a subset of these sequences identified several well-supported clades (Figure 1B). FaNaCs belong to a clade with members in several mollusk and annelids (not *Platynereis*) as well as one sequence from the brachiopod *Lingula anatina* and the phoronid *Phoronis australis*. This clade is sister to a clade containing *Platynereis* ENaC4-7 and related sequences from other annelids. A clade of nematode sequences is sister to FaNaC/ENaC4-7 and a clade of cephalochordate sequences is sister to all of these protostome sequences. There are also cnidarian sequences and other more distantly related protostome and deuterostome sequences, but the deep branching pattern of these was not resolved. The phylogenetic analysis thus revealed a bilaterian (possibly eumetazoan) family that includes both *Platynereis* ENaC4-7 and FaNaCs, with the two diverging sometime during the evolution of lophotrochozoans.

**Figure 1).**
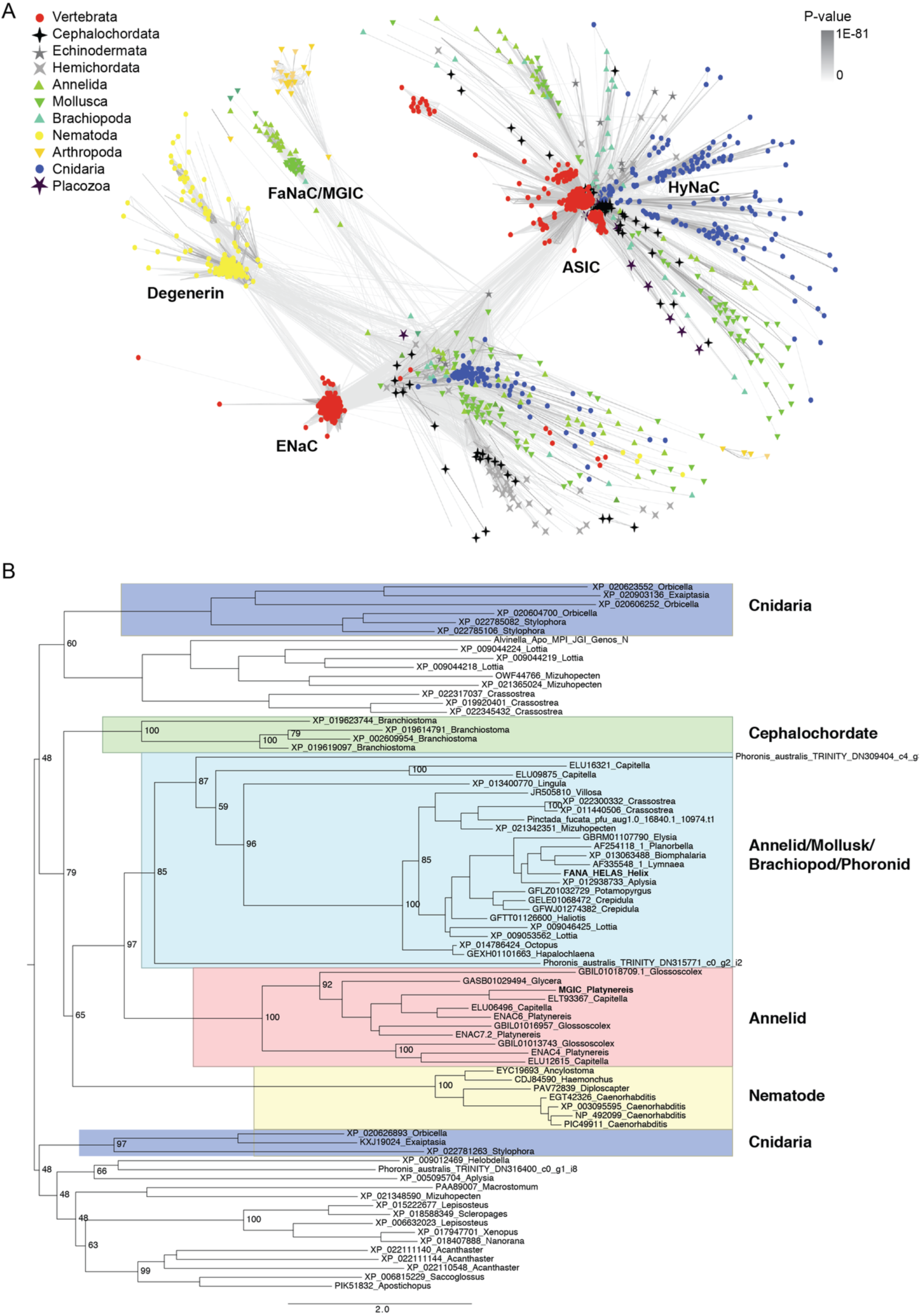
Phylogenetic position of *Platynereis* DEG/ENaCs. (**A**) Cluster map of DEG/ENaCs. Nodes correspond to different sequences and are colored based on taxonomy. Edges correspond to BLAST connections. (**B**) Maximum likelihood tree of FaNaCs and related sequences. Sequences are labeled by the GenBank identifier followed by the genus name. Well-resolved clades are shown with colored boxes. *Platynereis* MGIC and Helix FaNaC are shown in bold. Bootstrap values are shown for key branch points.

### A *Platynereis* DEG/ENaC subunit forms a myoinhibitory-peptide-gated ion channel

To express and functionally characterize the four *Platynereis* ENaCs, we injected the corresponding cRNAs in *Xenopus laevis* oocytes, either alone or in combination, and recorded whole cell currents. In healthy injected oocytes, we never observed increased basal current levels, suggesting that the channels are not constitutively active like ENaC or mouse BASIC (also named BLINaC) (25, 26). We therefore screened a library of 121 *Platynereis* neuropeptides for their effect on the four ENaCs from *Platynereis.* Initially, we divided the peptides into three pools (Supplementary Table 1). Pool 1 contained water soluble neuropeptides (63 peptides), excluding R/[F/Y]-amides, pool 2 contained R/[F/Y]-amides (12 peptides) and pool 3 contained peptides that required dimethylformamide as additional solvent (46 peptides). The three pools were supplemented to standard bath solution such that the concentration of the individual peptides was 1 µM. Strikingly, the first pool of peptides elicited robust currents of up to 2.8 µA in oocytes expressing ENaC5, but not in oocytes expressing ENaC4, 6 or 7. Currents appeared without delay upon application of the peptide mix and declined rapidly upon wash-out of the mix, suggesting that peptide pool 1 indeed contained a peptide that directly activated ENaC5. Pool 2 and 3 did not elicit currents with any ENaC subunit, excluding that ENaCs4-7 were activated by RFamides at 1 µM. It rather appeared that a peptide different from RF amides activated ENaC5. In the next step, we split active pool 1 in four subpools containing 15-16 peptides each. Two of these subpools elicited currents in oocytes expressing the ENaC5. As these two pools contained different peptides, this result suggested that two different peptides can activate ENaC5. These two subpools were then split again in final subpools containing 4 peptides each. Consistent with the previous round of screening, two of these four peptide-subpools elicited currents in oocytes expressing ENaC5. Finally, the eight peptides of these two subpools were screened individually and myoinhibitory peptides 2 and 7 (MIP2 and MIP7) were identified as peptides that elicited currents in oocytes expressing ENaC5.

**Table 1.** EC_50_ (M) values of the MIP GPCR tested with different MIP peptides. 95% confidence intervals for the EC_50_ (M) values are given.

| MIP1 | MIP2 | MIP3 | MIP4 | MIP5 | MIP6 | MIP7 | MIP8 | MIP9 | MIP10 | MIP11 |
| --- | --- | --- | --- | --- | --- | --- | --- | --- | --- | --- |
| $EC_{50}$ | 0,00001 | 0,000011 | 1,1E-08 | 5,5E-08 | 1,8E-08 | 6,6E-09 | 1,5E-08 | 0,000013 | 2,3E-08 | 0,000034 |
| 95% Confidence Intervals | 4.7e-6 to 2.2e-5 | 5.8e-6 to 2.2e-5 | 6.8e-9 to 2e-8 | 1.5e-8 to 1.9e-7 | 9.5e-9 to 3.5e-8 | 2.8e-9 to 1.5e-8 | 3.6e-9 to 6.4e-8 | 3.9e-6 to 4.3e-5 | 8.2e-9 to 6.7e-8 | 7.5e-7 to 0.0015 |

MIP2 and MIP7 are contained within the same proneuropeptide (12, 15) and share extensive sequence identity (Figure 2A), explaining why they both activated ENaC5. The independent identification of only MIP2 and MIP7 out of 121 peptides demonstrates the sensitivity and specificity of our screen. The *Platynereis* MIP proneuropeptide contains eleven MIP peptides (Figure 2A). As our neuropeptide library contained only MIP2 and MIP7, we tested all 11 MIPs (4 µM) for their ability to elicit currents in oocytes expressing ENaC5. Indeed, all MIPs elicited robust currents; we therefore named the *Platynereis* ENaC5 Myoinhibitory peptide-Gated Ion Channel (MGIC).

**Figure 2).**
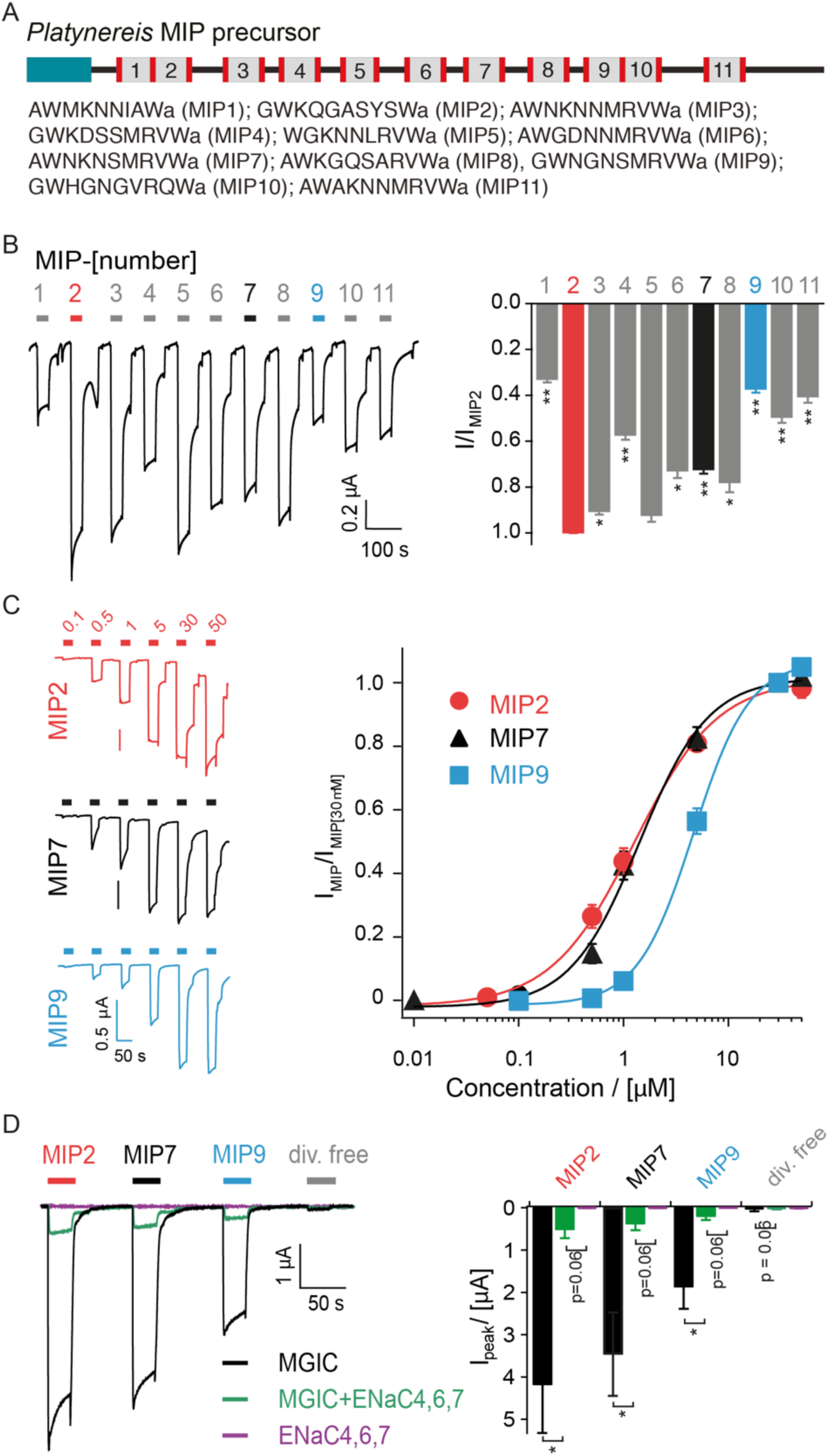
The DEG/ENaC ion channel MGIC is gated by myoinhibitory peptides. (**A**) Schematic of the *Platynereis* MIP proneuropeptide and sequence of the individual mature MIP peptides. Dibasic cleavage sites are shown in red. (**B**) Left, representative current trace of a MGIC-expressing oocyte activated by 4 µM of MIP1 – MIP11. Right, bar graph showing the mean ± SEM of MIP-gated currents normalized to the current gated by MIP2 (n = 8; *, p < 0.05 **, p < **0.01**, paired t test followed by Bonferroni correction). (**C**) Left, representative current traces of MGIC-expressing oocytes activated by increasing concentrations of MIP2, MIP7 or MIP9. Right, concentration response curves of the three MIPs normalized to the current evoked by 30 µM of the respective MIP. MIP2, n = 10–16; MIP7, n = 7–8; MIP9, n = 10–12. Error bars represent SEM. (**D**) Left, representative current traces of oocytes injected with different combinations of cRNA coding for DEG/ENaC subunits when exposed to 10 µM of MIP2, MIP7, MIP9 or divalent-free bath solution. Note that an equal amount of MGIC cRNA was injected for MGIC alone and for MGIC together with ENaC4, ENaC6 and ENaC7. Right, quantification of peak MGIC currents ± SEM activated by the indicated ligands (n = 6–7; *, p < 0.05, t test followed by Bonferroni correction).

The current amplitudes with different MIPs varied considerably (Figure 2B), with MIP2 inducing the largest and MIP1 inducing the smallest current amplitude (33±1% of the MIP2 gated current; n = 8; p <0.01, t-test with Bonferroni correction). For three representative MIPs, we determined the concentration-dependent activation of MGIC. We selected MIP2 and MIP7, which elicited large currents (MIP7: 73 ± 2% of the MIP2 gated current) and MIP9, which elicited relatively small currents (37 ± 1% of MIP2) at 4 µM (Figure 2B). All three MIPs activated MGIC in a concentration-dependent manner, with MIP2 being the most potent (Figure 2 C; EC_50_: 1.43 ± 0.22 µM, n = 16), followed by MIP7 (EC_50_: 1.73 ± 0.37 µM; n = 8) and MIP9 (EC_50_: 5.46 ± 0.77 µM; n = 12; t-test vs MIP2: p <0.001). Hill coefficients were in the range of 1.2 to 1.7 (MIP2: 1.18 ± 0.11; MIP7: 1.35 ± 0.06; MIP9: 1.72 ± 0.14), suggesting some cooperativity for MGIC gating by MIPs. As the potency of these three MIPs can explain the difference in the current amplitude when they were applied at a fixed concentration of 4 µM, this result suggests that the potency of MIPs at MGIC varies approximately tenfold from 1 to 10 µM. Among peptide gated DEG/ENaCs the affinity of MGIC for MIPs is comparable to the affinity of *Helix aspersa* FaNaC for FMRFamide (EC_50_: 2 µM; (5)). Affinities of the heteromeric HyNaCs for their peptide ligand Hydra-RFamide can be considerably higher (EC_50_ down to 40 nM), depending on their subunit composition (10). Comparing the affinity of MGIC for MIPs with the affinities of typical ionotropic receptors like AMPA or GABA_A_ receptors for their ligands, which is also in the low micromolar range (27, 28), MGIC can be classified as a high-affinity ionotropic receptor for MIPs.

In contrast to oocytes expressing MGIC, MIPs did not elicit a current in oocytes expressing ENaC4, 6 or 7. When cRNAs of ENaC4, 6 and 7 were co-injected with MGIC cRNA, currents gated by MIPs decreased relative to the currents when MGIC cRNA was injected alone (Figure 2D; I_MIP ave_: 0.37 ± 0.15 µA vs 3.2 ± 0.9 µA). Furthermore, oocytes co-injected with MGIC and with ENaCs4, 6 and 7 or injected with MGIC alone showed similar relative sensitivities towards the three MIPs tested, with MIP2 inducing the largest and MIP9 the smallest current (Figure 2D). These results suggest that MGIC did not form a heteromer with ENaC4, 6 or 7, but rather assembles into a functional homomeric channel upon expression in *Xenopus* oocytes. If the *Platynereis* ENaCs formed heteromers, we cannot distinguish their properties from a MGIC homomer based on the response to these three MIPs. This situation is similar to FaNaC from mollusks, which also assemble into functional homomers, but different from HyNaCs, which are obligate heteromers (10).

Next, we assessed the kinetics of MGIC currents gated by 10 µM MIP2 in more detail. The current rise time from 10% to 90% of the peak current was 0.68 ± 0.07 sec (n = 7), indicating a rapid activation of MGIC similar to HyNaC and FaNaC (5, 8). This rapid kinetics strongly suggests direct activation of MGIC by MIPs. MIP2-gated currents were partially (~25%) desensitizing with a mean time constant of 17.4 ± 3.4 sec (n = 7). This slow and partial desensitization is similar to FaNaC (5), but in contrast to HyNaCs which do not desensitize (9).

### Pore properties of MGIC

As DEG/ENaC ion channels usually have a high permeability for Na^+^ and a varying permeability for K^+^ ions, we first probed the dependence of MIP-gated currents on extracellular Na^+^ ions. When MGIC was activated by 1 µM MIP2 and Na^+^ was replaced by K^+^, the current amplitude dropped to 43 ± 4% of the current in the presence of Na^+^ (Figure 3A; n=10; t-test vs control, p=0.04). This result indicates that MGIC indeed conducts Na^+^ but is not strongly selective for this ion. The current remaining in the K^+^ containing bath could be mediated by K^+^ or divalent cations. To differentiate between these two possibilities, we activated MGIC and replaced Na^+^ by NMDG^+^, which is impermeable to most ion channels, but we kept the divalent cations in the bath solution. In this situation, MIP2 elicited a tiny sustained outward current (Figure 3B; n=8; t-test vs control, p=0.01). This is best explained by a K^+^ efflux, as divalent cations would produce inward currents at the holding potential of −70 mV. More importantly, a Ca^2+^ influx would cause the secondary activation of endogenous calcium-activated chloride channels (CaCCs), which would mediate large inward currents at −70 mV with a typical rapid kinetics (9) that we never observed for MGIC-expressing oocytes. To characterize the pore properties of MGIC in more detail, we recorded I-V curves in standard bath solution and in K^+^-containing bath solution (Figure 3 C). Both I-V curves were linear and had reversal potentials E_rev_ of 39.8 ± 1.5 mV (standard bath solution) and 1.4 ± 0.9 mV (K^+^ containing bath solution; n = 8; p < 0.001, t test), respectively. Based on these values a relative permeability P_Na+_/P_K+_ of 4.6 can be calculated. In conclusion, MGIC is a cation channel that is permeable to the monovalent cations Na^+^ and K^+^ but impermeable to divalent cations. MGIC shares relative permeability for monovalent cations with HyNaCs and impermeability to divalent cations with FaNaC, which shows a slightly higher sodium selectivity (P_Na+_ / P_K+_ > 10; (5)).

**Figure 3).**
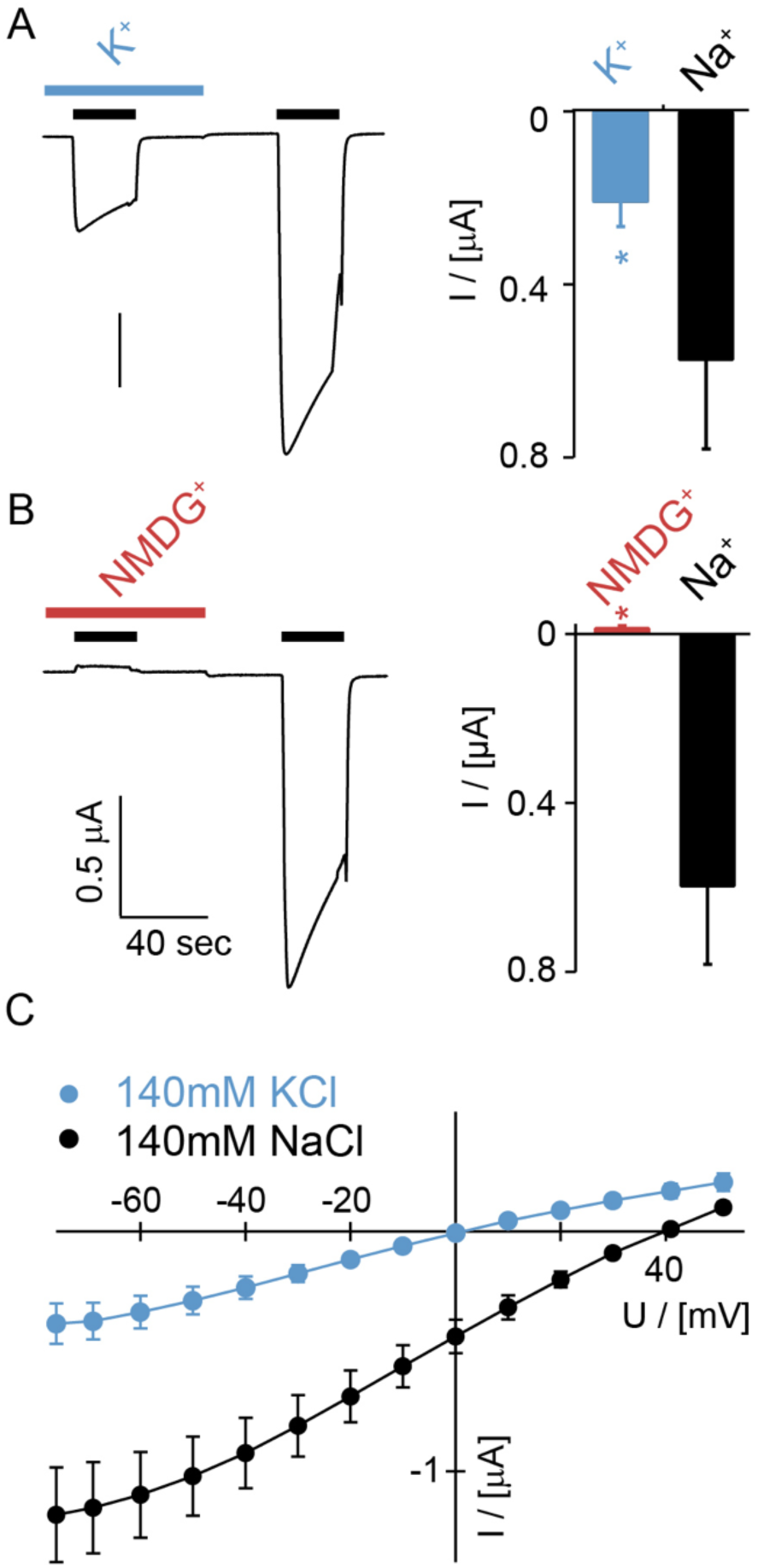
MGIC-5 conducts Na^+^ over K^+^. (**A** and **B**) Left, representative current traces of MGIC-expressing oocytes activated by 1 µM MIP2 (black bars) in bath solutions containing different ions. Bath solution with either K^+^ (A) or NMDG^+^ (B) replacing Na^+^ was present during the first activation by MIP2; during the second activation, standard Na^+^-containing bath solution was present. Right, quantification of the mean amplitudes ± SEM of MIP2-activated currents in the different bath solutions. n = 8–10; *, p < 0.05. (**C**) Current-voltage relationships of MGIC-expressing oocytes when activated by 1 µM MIP2 in the presence of bath solution containing Na^+^ or bath solution with K^+^ replacing Na^+^. Reversal potential in NaCl was 39.8 ± 1.5 mV and in KCl 1.4 ± 0.9 mV, PNa^+^/PK^+^ was 4.6. n = 8. Note that leakage currents in the absence of MIP2 were subtracted to reveal the MGIC-mediated current.

### Importance of tryptophan residues in MIP7

MIPs consist of nine to ten amino acid residues and contain two conserved tryptophans (Figure 2A). The first tryptophan is located at position 1 or 2 (W1 or W2) and the second at the C-terminal position (W9 or W10). We investigated the importance of these two tryptophans for MGIC activation by changing them into alanine in MIP7.

When W2 was mutated into alanine (MIP7-W2A) the current amplitude induced by 10 µM of this peptide decreased to 12 ± 1% relative to 10 µM MIP7 (Figure 4A; n = 7; p < 0.01, t test with Bonferroni correction). Changing W10 to alanine (MIP7-W10A) abolished activation of MGIC (Figure 4A; 0.3 ± 0.1% relative to MIP7; n = 7; p < 0.01). Changing W2 and W10 together also abolished activation of MGIC (Figure 4A; 0.1 ± 0.1% relative to MIP7; n = 7; p < 0.01).

**Figure 4).**
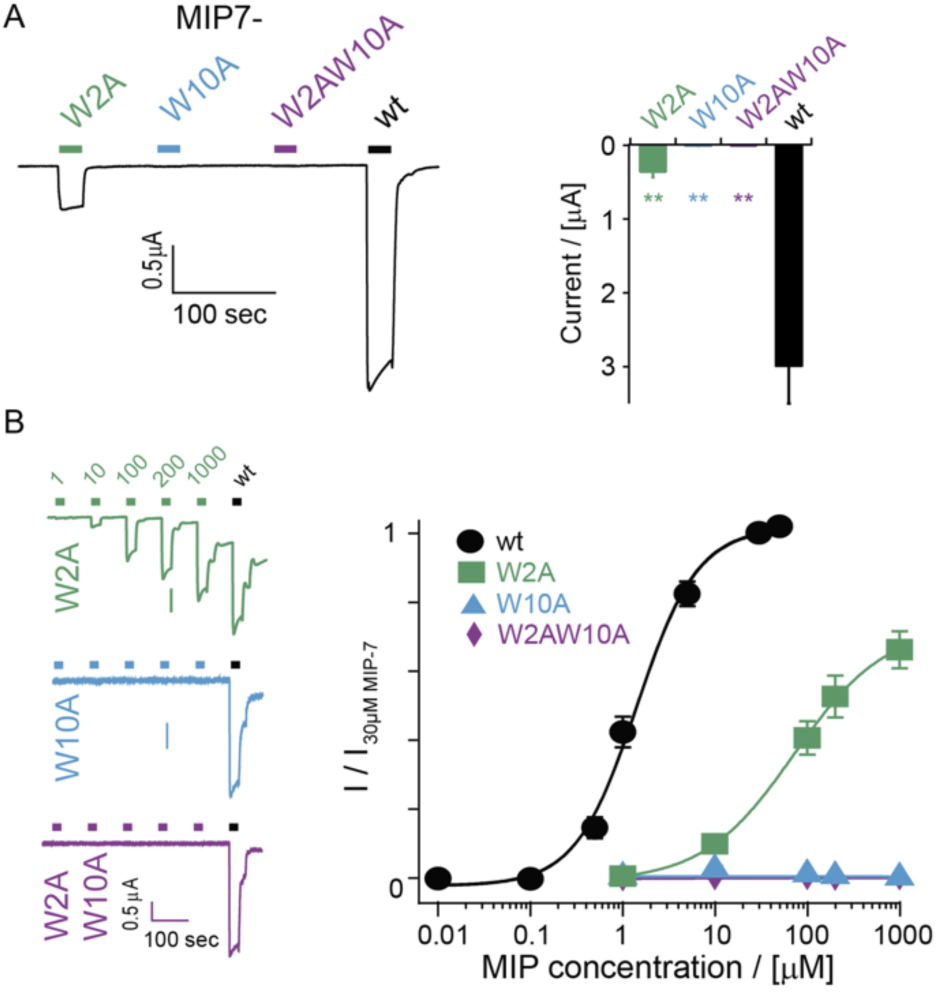
The C-terminal tryptophan is vital for MIP gating of MGIC. (**A**) Left, representative current trace of a MGIC-expressing oocyte exposed to 10 µM of MIP7 or three MIP7 variants containing individual amino acid substitutions. Right, bar graph showing the mean current amplitudes ± sem. **, p < 0.01, Bonferroni; n = 7. (**B**) Left, representative current traces of MGIC-expressing oocytes exposed to increasing concentrations of MIP7 mutants; as a reference, 30 µM MIP7 was applied at the end of each experiment. Right, concentration response curves of MIP7 and its variants. Data for MIP7 wt taken from Figure 2 normalized to the current evoked by 30 µM MIP7. n = 8; error bars represent sem.

We recorded concentration-response curves for the three MIP7 variants and compared them to the MIP7 concentration-response curve. The W2A substitution strongly reduced the apparent affinity (Figure 4B; cf. Figure 2D; EC_50_: 80 ± 14 µM; n = 8; p < 0.001, t test). Substitutions W10A and W2AW10A did not produce reliable currents even at a peptide concentration of 1 mM. Overall both tryptophans seem to be important for MGIC activation, with the C-terminal tryptophan being essential.

### Common DEG/ENaC blockers inhibit MGIC

A common feature among DEG/ENaCs is their sensitivity to the inhibitor amiloride and related substances. To determine the pharmacological profile of MGIC, we tested the inhibition by amiloride, benzamil, and phenamil. We also tested inhibition by the diarylamidine diminazene, which inhibits ASICs, BASIC and HyNaCs but not ENaC (10, 29, 30). At 50 µM, all four drugs reduced the peak currents gated by 1 µM MIP2 with the following rank order: diminazene > amiloride > phenamil > benzamil (Figure 5A; diminazene: 8 ± 1% of control; amiloride: 25 ± 1%; p <0.001; benzamil: 74 ± 3%; p <0.005; phenamil 45 ± 4%; p < 0.001; n = 7; t test with Bonferroni correction).

**Figure 5).**
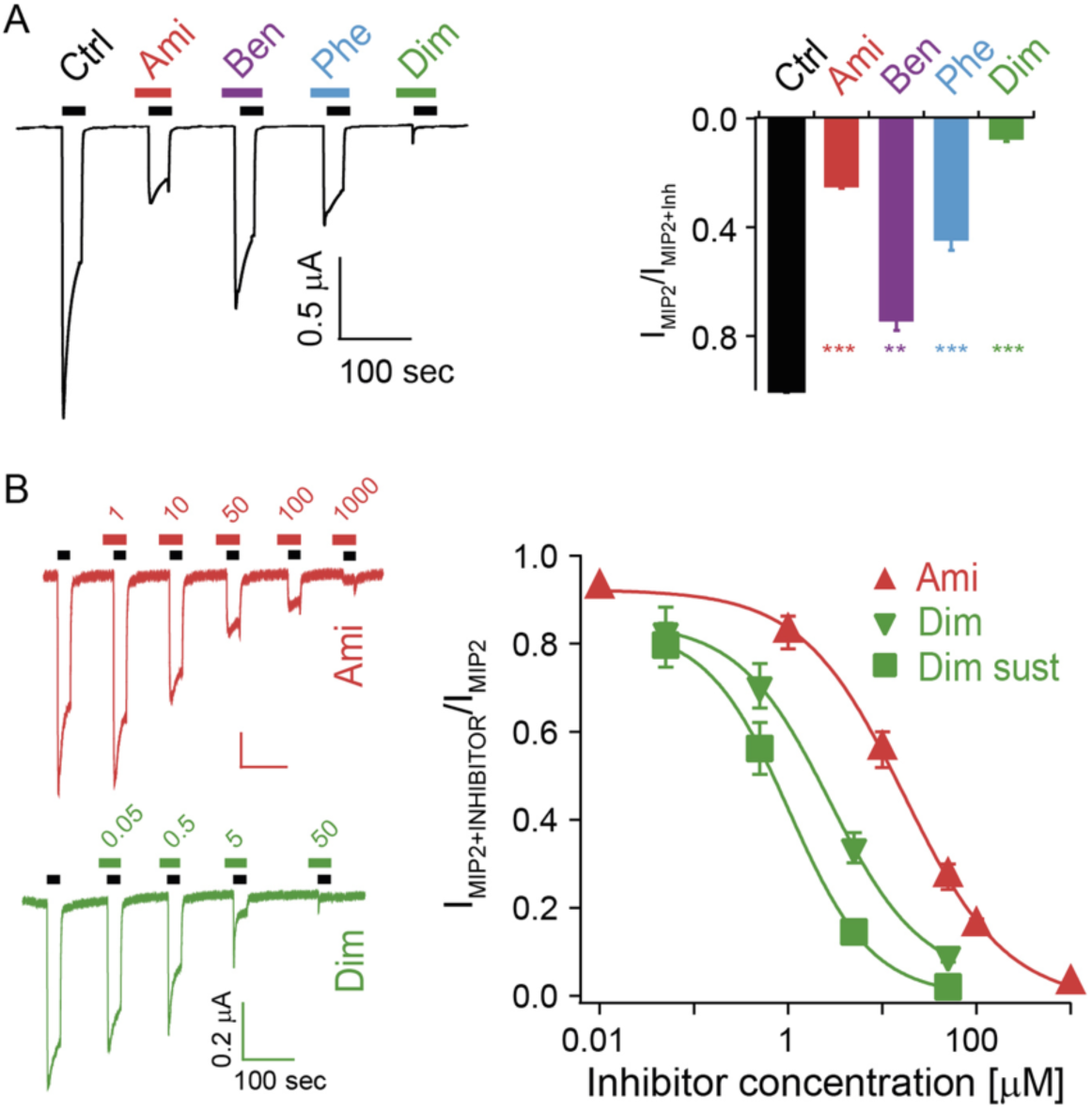
MGIC is blocked by typical DEG/ENaC inhibitors. (**A**) Left, representative trace of a MGIC-expressing oocyte activated by 1 µM MIP2 alone or in the presence of 50 µM amiloride (Ami), benzamil (Ben), phenamil (Phe) or diminazene (Dim). Right, bar graph illustrating mean current amplitudes ± SEM normalized to the MIP2-activated current in the absence of inhibitors. n = 7; **, p < 0.01; ***, p < 0.001 test with Bonferroni correction. (**B**) Left, representative current traces showing the response to 1 µM MIP2 when amiloride or diminazene were pre- and coapplied (concentrations in µM). Note the effect of diminazene on the current kinetics and the washout effect upon amiloride/MIP2 removal. Right, quantification of the currents normalized to the current of 1 µM MIP2 in the absence of inhibitors (amiloride, n = 5 for 0.01 µM and n = 12–13 for all other concentrations; diminazene, n = 8). In the case of diminazene, a quantification of the sustained current (Dim sust) revealed an even higher apparent affinity of MGIC to diminazene.

We determined complete concentration-response curves for diminazene and amiloride, the two most potent inhibitors. At high concentrations (> 1 µM) currents in the presence of diminazene had a biphasic appearance: the peak amplitude was reduced but the inhibitor further accelerated the current decay of MGIC. Such a behavior is expected when diminazene is a relatively slow pore blocker of MGIC (31). Therefore, we quantified the reduction of the amplitude by diminazene both at the beginning of a current deflection (quick but partial inhibition) and after full inhibition, yielding an IC_50_ of 2.4 ± 0.6 µM for quick, partial inhibition and an IC_50_ of 0.73 ± 0.16 µM for full inhibition (Figure 5B; n = 7–8). Amiloride reduced currents with an apparent IC_50_ of 13 ± 2 µM (Figure 5B; n = 12). Thus, the apparent affinity of MGIC for diminazene and amiloride is similar to ASICs (29) and BASIC (26, 30). HyNaCs also have a high affinity for diminazene (10) but a >10-fold lower affinity for amiloride and its analogs (8). In contrast the lower potency of benzamil and phenamil compared to amiloride are distinct from HyNaCs, ASICs as well as ENaC (8, 32, 33).

### Different MIP peptides activate MAG and MGIC with different potencies

*Platynereis* MIP7 was previously shown to signal by a G-protein coupled receptor (15). This MIP-Activated GPCR (MAG) is the ortholog of the *Drosophila* MIP GPCR, the sex peptide receptor (34). To further characterize the role of MAG in mediating MIP signals, we recorded dose-response curves for all 11 MIP peptides (Figure 6).

**Figure 6).**
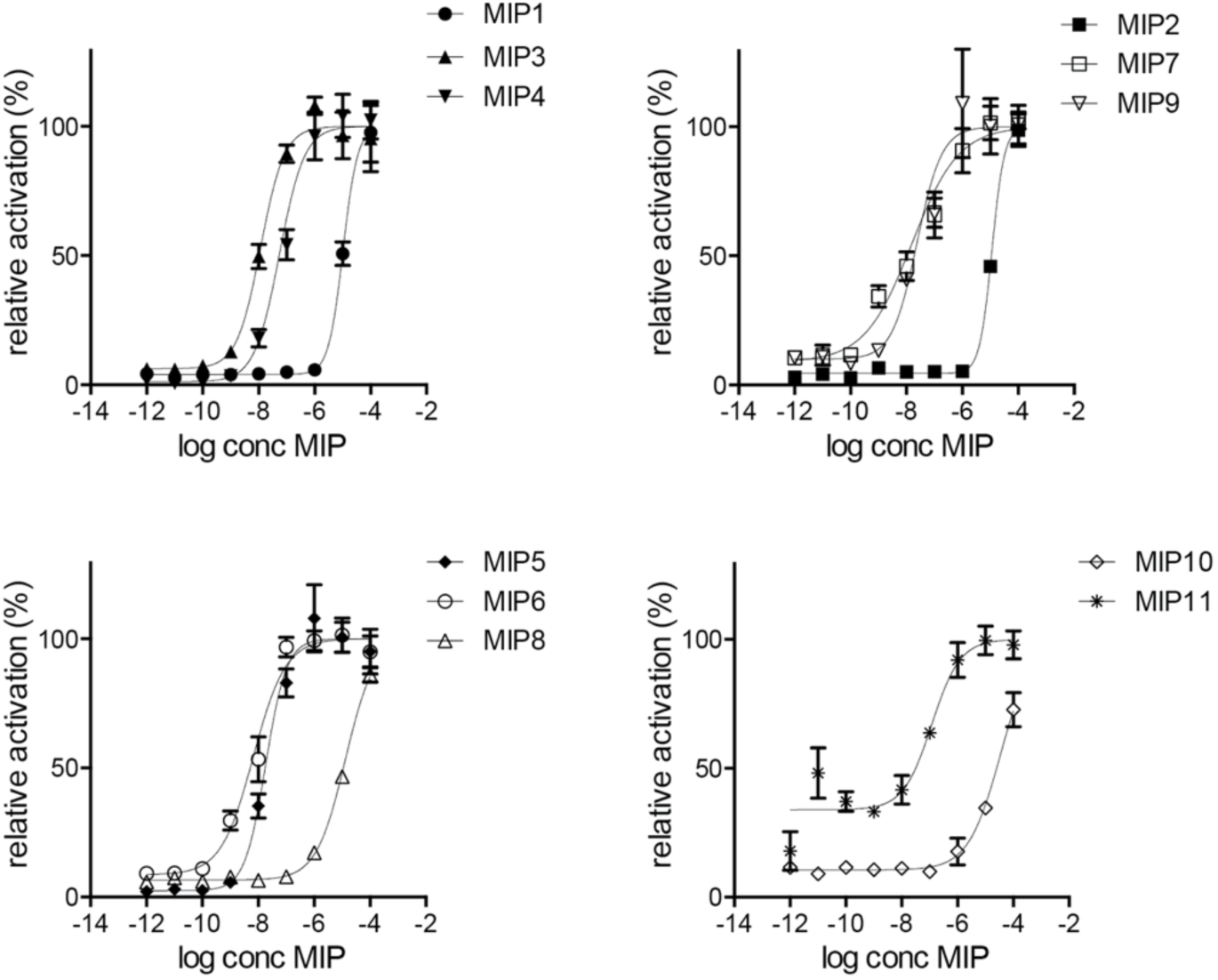
Individual dose-response curves of *Platynereis* MAG treated with varying concentrations of MIP peptides. Data, representing luminescence units relative to the maximum of the fitted dose-response curves, are shown as mean ± SEM (n = 3). EC_50_ values and significance values are listed in Table 1.

We found that MIP3, MIP4, MIP5, MIP6, MIP7, MIP9 and MIP11 activated MAG with EC_50_ values in the nanomolar range, with MIP6 being the most potent ligand (Figure 6 and Table 1). In contrast, MIP1 (AWamide), MIP2 (SWamide), MIP8 (VWamide) and MIP10 (QWamide) were poor ligands, activating the receptor only at micromolar concentrations (Figure 6 and Table 1).

### Differential expression of MAG and MGIC in *Platynereis* larvae

Next we wanted to test if MAG and MGIC participate in different signaling networks in *Platynereis*. We thus mapped the expression pattern of MGIC to a virtual larval brain of spatially mapped single-cell transcriptome data (17, 35) (Figure 7A). We have previously mapped the expression of MAG and MIP and established peptidergic signaling networks between the expressing cells by treating MIP-expressing cells as source nodes and MAG-expressing cells as target nodes (17)(Figure 7B). We also mapped the connectivity between MIP-expressing and MGIC-expressing cells (Figure 7 C). This analysis revealed that MAG and MGIC are expressed in largely non-overlapping sets of cells and form independent signaling networks. MAG-expressing cells are found in the neurosecretory anterior nervous system (15, 17) whereas MGIC is more broadly expressed in the larval episphere. This analysis suggests that MIP and MGIC participate in different signaling networks with little overlap in the nervous system of *Platynerei*s.

**Figure 7).**
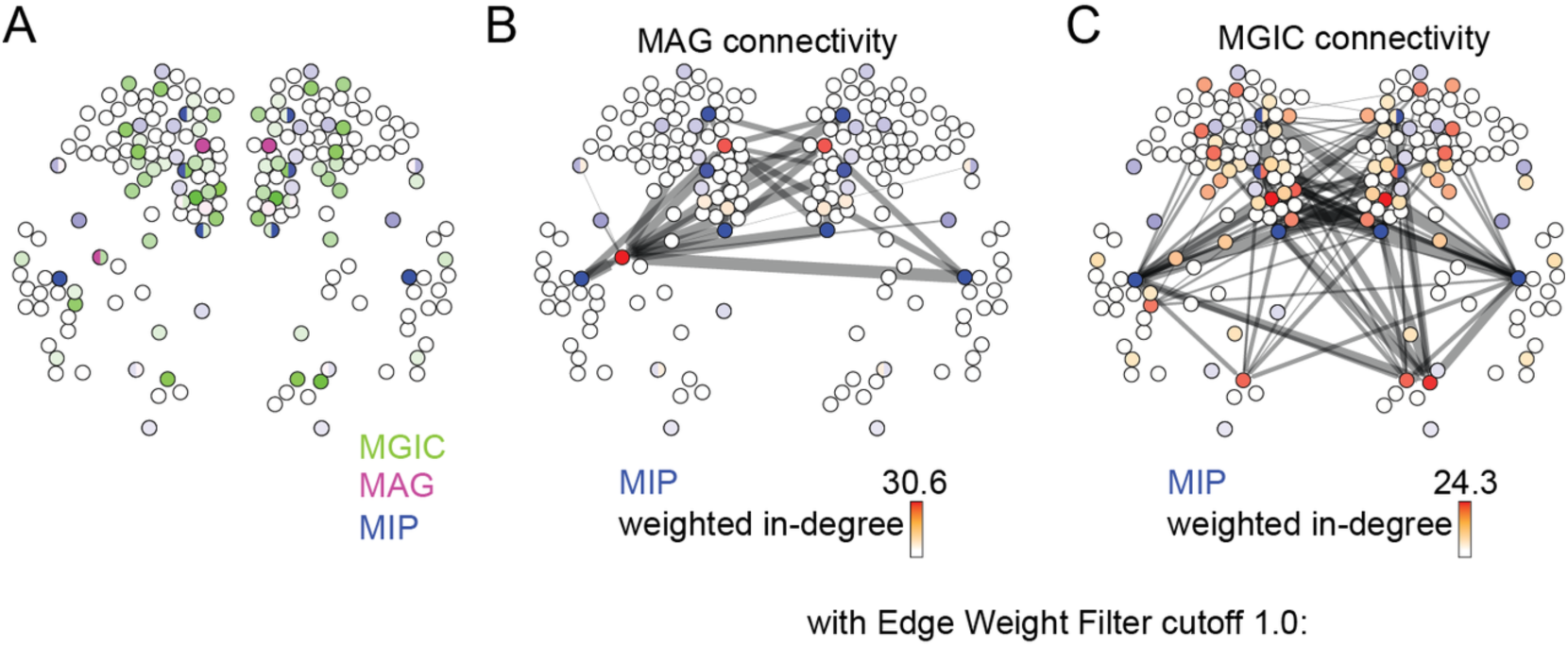
Mapping the expression of MIP, MAG and MGIC to spatially mapped single cell transcriptomes in the *Platynereis* larval head. (**A**) Expression of MIP, MAG and MGIC projected on the single-cell map. Color intensity of nodes reflects magnitude of normalized log10 gene expression. (**B**) Connectivity map of MIP and MAG, colored by weighted in-degree (red) and proneuropeptide log10 normalized expression (blue). Line thickness was determined by the geometric mean of log10 normalized proneuropeptide expression of signaling cell and log10 normalized receptor expression of the corresponding receiving cell. (**C**) Connectivity map of MIP and MGIC, colored by weighted in-degree (red) and proneuropeptide log10 normalized expression (blue). Line thickness was determined by the geometric mean of log10 normalized proneuropeptide expression of signaling cell and log10 normalized receptor expression of the corresponding receiving cell.

## Discussion

*Platynereis* MGIC represents the first ion channel that is gated by a neuropeptide that is not an RFamide. Wamides belong to an ancient neuropeptide family present in protostomes, cnidarians and the placozoan *Trichoplax adhaerens* but are absent from deuterostomes (15, 36, 37). The orthology group of MGIC includes only annelid sequences, but the divergence of this clade must predate the divergence of annelids, mollusks, phoronids and brachiopods, given that the FaNaC sister group contains representatives from these phyla. Our phylogenetic analysis also uncovered a clade of nematode, amphioxus and cnidarian sequences that are related to MGIC/FaNaC peptide gated channels. Further experimental analysis of a larger number of channels will be needed to reconstruct the patterns of ligand evolution in this family. The discovery of a Wamide-gated channel suggests a larger ligand diversity of peptide-gated channels. The phylogeny also suggests clade-specific duplications (four MGIC paralogs in *Platynereis)* and frequent losses (no MGIC ortholog was yet found in mollusks). It is also possible that some of the *Platynereis* MGIC paralogs (ENaC4, ENaC6, and ENaC7) have lost the property of peptide-gating since they were not activated by any of the 121 neuropeptides tested. An alternative explanation would be that ENAC4, 6 and 7 form heteromeric peptide-gated channels with other uncharacterized DEG/ENaC subunits in *Platynereis.* An exciting future avenue will be to use a broader sampling of lophotrochozoan taxa and reconstruct the evolution of ligand specificities of FaNaC/MGIC channels.

An interesting property of the MIP system in *Platynereis* is the possibility of dual signaling by MGIC and MAG to distinct target neurons. In *Platynereis,* MIP peptides induce larval settlement by inhibiting ciliary swimming and are involved in the initiation of larval feeding following settlement (15, 16). These different effects may be mediated by either ionotropic or metabotropic signals. The effect of MIP on ciliary beating in *Platynereis* larvae is at least partly mediated by MAG, given that morpholino knock-down of this GPCR abrogates the effect of MIP on ciliary closures (15). Further knock-down or knockout studies on both MGIC and MAG will be needed to dissect the contributions of these two receptors to MIP signaling.

Dual signaling is likely also a characteristic *of Aplysia californica* and *Helix* FMRFamide neuropeptides. *In Aplysia* and probably other mollusks the FMRFamide-gated FaNaC coexists with a GPCR (XP005110250) that is orthologous to an FMRFamide G-protein coupled receptor from *Platynereis* and the brachiopod *Terebratalia transversa* (18, 38). It is thus possible that divergent peptidergic signaling through both ionotropic and metabotropic receptors is a more general feature of peptidergic signaling systems.

The presence of a dual mode of signaling will also have consequences for the evolution of the shared neuropeptide ligands. In proneuropeptides containing multiple copies of similar peptides, it is expected that the ligand specificities will diverge during evolution and different copies of the peptide will be tuned to one or the other receptor. A similar evolutionary process has likely happened in the *Platynereis* MIP precursor. For example, MIP2 is a poor ligand of MAG but is the most potent ligand of MGIC. MIP2 is an SWamide, a unique dipeptide ending out of the 11 MIPs. Such a process of evolutionary subfunctionalization may contribute to maintaining peptide diversity and copy number within the same precursor.

In summary, our results suggest that peptide-gated channels are widespread in lophotrochozoans and that their peptide ligands belong to different neuropeptide families.

## Materials and Methods

### Cloning of ENaCs and cRNA synthesis

We identified four *Platynereis* DEG/ENaC channels based on high sequence similarity to the *Helix aspersa* FaNaC receptor. The sequences were cloned from a *Platynereis* mixed-stages cDNA library by PCR. A restriction site and the Kozak sequence were introduced before the start codon into the forward primers. The primer sequences are: Pdu_ENaC4 fw ACAATAGGATCCCGCCACCATGGCGCATCCGCTACACCGG; Pdu_ ENaC4 rv ACAATAGGTACCTCATACCGCTGTCAGTTTGTCACCTACTTCAAAG; Pdu_ ENaC5 fw ACAATAGGATCCCGCCACCATGGCGCTACGTGCACTGATGCAG; Pdu_ ENaC 5 rv ACAATAGGTACCTCACACCATATGTGAAGGGTCCATGGG; Pdu_ ENaC 6 fw ACAATACTCGAGCGCCACCATGATGGTCCTCACCCTCCTTTTGAACTTG; Pdu_ ENaC 6 rv ACAATAACCGGTTCACAAAGTTGTGTTTTCCATTCTAAACTGAAGGTC; Pdu_ ENaC 7 fw ACAATAGGATCCCGCCACCATGGCTGGCAAAAGGGTCCTAAAATCG; Pdu_ ENaC 7 rv ACAATAGGTACCTTATCCAAGGATTGCTGAAGAGCATTCTAC. The *Platynereis* ENaC channel cDNA sequences were submitted to Genbank under the following names and accession numbers: ENaC4, MG844425; MGIC, MG844426; ENaC6, MG844427; ENaC7, MG844428. For the functional expression in *Xenopus laevis* oocytes, the cDNAs were subcloned into the pRSSP oocyte expression vector (20). This vector contains the 5’-untranslated region of the *Xenopus* β-globin and a poly(A) tail to improve protein expression and RNA stability. After cloning, the inserts were verified by sequencing. For *in vitro* cRNA synthesis, vectors were first linearized by restriction digestion. This linearized vector served as template for the in vitro cRNA synthesis using SP6 RNA Polymerase to yield capped cRNA (mMessage mMaschine kit; Ambion, Austin, TX, USA).

### Phylogenetic analysis and expression mapping

DEG/ENaC sequences were collected at the NCBI database. We first used psi-blast with two iterations in the NCBI non-redundant, expressed sequence tag (EST) and transcriptome shotgun assembly (TSA) databases using *Helix* FaNaC as a query. Closely related sequences were removed with the program CD-HIT (21) at an identity cutoff of 90% resulting in 3773 sequences. For the initial filtering of sequence, similarity-based cluster maps were generated with CLANS2 (22) using the BLOSUM62 matrix and a P-value cutoff of 1e-50. We selected a cluster of sequences including ENaCs, nematode Degenerins, FaNaC/MGIC and ASICs. This subset was re-clustered and is shown in Figure 1A. For phylogenetic tree reconstruction, we collected a smaller set of sequences by blasting with *Helix* FaNaC. We aligned the protein sequences with the program MUSCLE (23). We trimmed the poorly aligned regions with the program TrimAI (24) in “Automated 1” mode. Maximum likelihood trees were calculated with IQ-TREE using automatic model selection (http://www.iqtree.org/). The best-fitting model was the WAG+F+R8 model. Trees were visualised with FigTree (http://tree.bio.ed.ac.uk/software/figtree/).

The expression patterns of MGIC were projected to the spatially mapped single-cell transcriptome data as previously described (17).

### Electrophysiology

Animal care and surgery of *Xenopus laevis* frogs was conducted according to protocols approved by the state office for nature, environment and consumer protection (LANUV) of the state North Rhine-Westphalia (NRW). To obtain individual oocytes, ovaries of anaesthetized female frogs were removed by surgery, manually dissected and subsequently treated with collagenase type 2 (Worthington Biochemical Corporation, Lakewood, NJ, USA) for 2 hours. Oocytes of stage V or VI were selected and injected with 2–8 ng of cRNA coding for MGIC or ENaC4, 6 or 7. The cRNA injected oocytes were kept for 2–5 days at 19°C in OR-2 medium containing (in mM): 82.5 NaCl, 2.5 KCl, 1.0 Na_2_HPO_4_, 5.0 HEPES, 1.0 MgCl_2_, 1.0 CaCl_2_, 0.5 g/liter polyvinylpyrrolidone, 1,000 U/l penicillin and 10 mg/l streptomycin; pH was adjusted to 7.3.

Whole cell currents of manually defolliculated *Xenopus* oocytes were measured at ambient temperature with a two-electrode voltage clamp (TEVC) system. The system uses a Turbo Tec-03X TEVC amplifier (npi electronics, Tamm, Germany) for the measurement of whole cell currents and a programmable pipetting workstation for fast solution exchange (ScreeningTool; npi electronic). Micropipettes with a resistance of 1–3 MOhm filled with 3 mM KCl served as intracellular electrodes. Current traces were recorded at a sampling rate of 100 Hz (voltage steps) and 500 Hz (all other recordings) with the software CellWorks 6.2.1 (npi). Cells were clamped at a holding potential of −70 mV except in experiments examining the current-voltage relationship of ionic currents; in this case the holding potential was changed in a step-like fashion.

If not stated otherwise, standard bath solution contained (in mM): 140 NaCl, 1.8 CaCl_2_, 1 MgCl_2_, 10 HEPES. Peptides and inhibitors were supplemented to the standard bath solution as indicated. For ion exchange experiments bath solutions with Na^+^ replaced by K^+^ or NMDG^+^ were used. The divalent free solution contained (in mM) 140 NaCl, 0.79 CaCl_2_, 2 EGTA, 10 HEPES and 0.1 flufenamic acid to yield residual free Ca^2+^ concentration of 10 nM. For all solutions pH was adjusted to 7.4 with NaOH, KOH or HCl as appropriate.

Synthetic peptides were obtained from Genscript (Nanjing, China). All other chemicals used for electrophysiology were obtained from Carl Roth (Karlsruhe, Germany), Sigma-Aldrich (St. Louis, Missouri, USA) or VWR (Radnor, Pennsylvania, USA).

### Data analysis

Current traces were visualized and analyzed with the software CellWorks Reader 6.2.2 (npi). Data analysis was performed using Microsoft Excel and Igor Pro (Vesion 5.02, WaveMetrics, Lake Oswego, OR, USA). Dose response curves were fit with a Hill function:

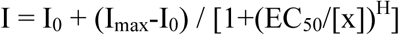

where I represents the current, [x] the concentration of the considered substance x, I_0_ and I_max_ the minimal and maximal current, respectively, EC_50_ the concentration, at which 50% of the maximal current is reached, and H the Hill coefficient.

I-V curves were obtained by subtracting currents in the absence of agonist from currents in its presence to correct for background currents. To obtain reversal potentials, we used a linear fit of the three adjacent data points between which current reversed its sign.

To obtain P_Na+_ / P_K+_, the shift in reversal potentials (ΔE_rev_) when Na^+^ was replaced by K^+^ in the bath solution was determined. Assuming constant intracellular concentrations of the considered ions, P_Na+_ / P_K+_ was then calculated with an equation derived from the Goldman–Hodgkin–Katz equation:

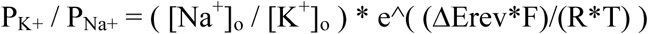

Where F represents the Faraday constant, R the gas constant, and T the absolute temperature.

Results are reported as mean ± s.e.m., error bars in figures also represent s.e.m. Paired or unpaired Student’s t-test (t-test) was used to test for significance when comparing two groups. For comparing multiple groups, p values obtained by t-tests were multiplied by the number of tests conducted on the same dataset (Bonferroni correction). For all measurements, oocytes of at least two different frogs were used.

## Acknowledgements

PB was supported by the International Max Planck Research School (IMPRS) “From Molecules to Organisms.” EAW was supported by a grant from the DFG – Deutsche Forschungsgemeinschaft (Reference no. JE 777/1).

## Author Contributions

AS, PB, KA, EW, SG and GJ designed and performed research and analyzed data. AS, GJ and SG wrote the paper.

## References

1. Nusbaum, M.P., Blitz, D.M. and Marder, E. (2017) Functional consequences of neuropeptide and small-molecule co-transmission. Nature Reviews. Neuroscience, 18, 389–403.

2. Jékely, G., Melzer, S., Beets, I., Grunwald Kadow, I.C., Koene, J., Haddad, S., et al. (2018) The long and the short of it – a perspective on peptidergicregulation of circuits and behaviour. The Journal of Experimental Biology, 2018.

3. Krishnan, A. and Schiöth, H.B. (2015) The role of G protein-coupled receptors in the early evolution of neurotransmission and the nervous system. The Journal of Experimental Biology, 218, 562–571.

4. Cottrell, G.A., Green, K.A. and Davies, N.W. (1990) The neuropeptide Phe-Met-Arg-Phe-NH2 (FMRFamide) can activate a ligand-gated ion channel in *Helix* neurones. Pflugers Archiv: European Journal of Physiology, 416, 612–614.

5. Lingueglia, E., Champigny, G., Lazdunski, M. and Barbry, P. (1995) Cloning of the amiloride-sensitive FMRFamide peptide-gated sodium channel. Nature, 378, 730–733.

6. Furukawa, Y., Miyawaki, Y. and Abe, G. (2006) Molecular cloning and functional characterization of the *Aplysia* FMRFamide-gated Na+ channel. Pflugers Archiv: European Journal of Physiology, 451, 646–656.

7. Golubovic, A., Kuhn, A., Williamson, M., Kalbacher, H., Holstein, T.W., Grimmelikhuijzen, C.J.P., et al. (2007) A peptide-gated ion channel from the freshwater polyp *Hydra*. The Journal of Biological Chemistry, 282, 35098–35103.

8. Dürrnagel, S., Kuhn, A., Tsiairis, C.D., Williamson, M., Kalbacher, H., Grimmelikhuijzen, C.J.P., et al. (2010) Three homologous subunits form a high affinity peptide-gated ion channel in *Hydra*. The Journal of Biological Chemistry, 285, 11958–11965.

9. Dürrnagel, S., Falkenburger, B.H. and Gründer, S. (2012) High Ca(2+) permeability of a peptide-gated DEG/ENaC from *Hydra*. The Journal of General Physiology, 140, 391–402.

10. Assmann, M., Kuhn, A., Dürrnagel, S., Holstein, T.W. and Gründer, S. (2014) The comprehensive analysis of DEG/ENaC subunits in *Hydra* reveals a large variety of peptide-gated channels, potentially involved in neuromuscular transmission. BMC Biology, 12, 84.

11. Gründer, S. and Assmann, M. (2015) Peptide-gated ion channels and the simple nervous system of *Hydra*. The Journal of Experimental Biology, 218, 551–561.

12. Conzelmann, M., Williams, E.A., Krug, K., Franz-Wachtel, M., Macek, B. and Jékely, G. (2013) The neuropeptide complement of the marine annelid *Platynereis dumerilii*. BMC Genomics, 14, 906.

13. Tessmar-Raible, K., Raible, F., Christodoulou, F., Guy, K., Rembold, M., Hausen, H., et al. (2007) Conserved sensory-neurosecretory cell types in annelid and fish forebrain: insights into hypothalamus evolution. Cell, 129, 1389–1400.

14. Conzelmann, M., Offenburger, S.-L., Asadulina, A., Keller, T., Münch, T.A. and Jékely, G. (2011) Neuropeptides regulate swimming depth of *Platynereis* larvae. Proceedings of the National Academy of Sciences of the United States of America, 108, E1174–83.

15. Conzelmann, M., Williams, E.A., Tunaru, S., Randel, N., Shahidi, R., Asadulina, A., et al. (2013) Conserved MIP receptor-ligand pair regulates *Platynereis* larval settlement. Proceedings of the National Academy of Sciences of the United States of America, 110, 8224–8229.

16. Williams, E.A., Conzelmann, M. and Jékely, G. (2015) Myoinhibitory peptide regulates feeding in the marine annelid *Platynereis*. Frontiers in zoology, 12, 1.

17. Williams, E.A., Verasztó, C., Jasek, S., Conzelmann, M., Shahidi, R., Bauknecht, P., et al. (2017) Synaptic and peptidergic connectome of a neurosecretory center in the annelid brain. eLife, 6.

18. Bauknecht, P. and Jékely, G. (2015) Large-Scale Combinatorial Deorphanization of *Platynereis* Neuropeptide GPCRs. Cell reports, 12, 684–693.

19. Bauknecht, P. and Jékely, G. (2017) Ancient coexistence of norepinephrine, tyramine, and octopamine signaling in bilaterians. BMC Biology, 15, 6.

20. Bässler, E.L., Ngo-Anh, T.J., Geisler, H.S., Ruppersberg, J.P. and Gründer, S. (2001) Molecular and functional characterization of acid-sensing ion channel (ASIC) 1b. The Journal of Biological Chemistry, 276, 33782–33787.

21. Li, W. and Godzik, A. (2006) Cd-hit: a fast program for clustering and comparing large sets of protein or nucleotide sequences. Bioinformatics, 22, 1658–1659.

22. Frickey, T. and Lupas, A. (2004) CLANS: a Java application for visualizing protein families based on pairwise similarity. Bioinformatics, 20, 3702–3704.

23. Edgar, R.C. (2004) MUSCLE: a multiple sequence alignment method with reduced time and space complexity. BMC Bioinformatics, 5, 113.

24. Capella-Gutiérrez, S., Silla-Martínez, J.M. and Gabaldón, T. (2009) trimAl: a tool for automated alignment trimming in large-scale phylogenetic analyses. Bioinformatics, 25, 1972–1973.

25. Canessa, CM., Schild, L., Buell, G., Thorens, B., Gautschi, I., Horisberger, J.D., et al. (1994) Amiloride-sensitive epithelial Na+ channel is made of three homologous subunits. Nature, 367, 463–467.

26. Wiemuth, D. and Gründer, S. (2010) A single amino acid tunes Ca2+ inhibition of brain liver intestine Na+ channel (BLINaC). The Journal of Biological Chemistry, 285, 30404–30410.

27. Hollmann, M. and Heinemann, S. (1994) Cloned glutamate receptors. Annual Review of Neuroscience, 17, 31–108.

28. Sigel, E., Baur, R., Trube, G., Möhler, H. and Malherbe, P. (1990) The effect of subunit composition of rat brain GABAA receptors on channel function. Neuron, 5, 703–711.

29. Chen, X., Qiu, L., Li, M., Dürrnagel, S., Orser, B.A., Xiong, Z.-G., et al. (2010) Diarylamidines: high potency inhibitors of acid-sensing ion channels. Neuropharmacology, 58, 1045–1053.

30. Wiemuth, D. and Gründer, S. (2011) The pharmacological profile of brain liver intestine Na+ channel: inhibition by diarylamidines and activation by fenamates. Molecular Pharmacology, 80, 911–919.

31. Schmidt, A., Rossetti, G., Joussen, S. and Gründer, S. (2017) Diminazene Is a Slow Pore Blocker of Acid-Sensing Ion Channel 1a (ASIC1a). Molecular Pharmacology, 92, 665–675.

32. Leng, T.-D., Si, H.-F., Li, J., Yang, T., Zhu, M., Wang, B., et al. (2016) Amiloride analogs as asic1a inhibitors. CNS Neuroscience & Therapeutics, 22, 468–476.

33. Kleyman, T.R. and Cragoe, E.J. (1988) Amiloride and its analogs as tools in the study of ion transport. The Journal of Membrane Biology, 105, 1–21.

34. Kim, Y.-J., Bartalska, K., Audsley, N., Yamanaka, N., Yapici, N., Lee, J.-Y., et al. (2010) MIPs are ancestral ligands for the sex peptide receptor. Proceedings of the National Academy of Sciences of the United States of America, 107, 6520–6525.

35. Achim, K., Pettit, J.-B., Saraiva, L.R., Gavriouchkina, D., Larsson, T., Arendt, D., et al. (2015) High-throughput spatial mapping of single-cell RNA-seq data to tissue of origin. Nature Biotechnology, 33, 503–509.

36. Jékely, G. (2013) Global view of the evolution and diversity of metazoan neuropeptide signaling. Proceedings of the National Academy of Sciences of the United States of America, 110, 8702–8707.

37. Nikitin, M. (2015) Bioinformatic prediction of *Trichoplax adhaerens* regulatory peptides. General and Comparative Endocrinology, 212, 145–155.

38. Thiel, D., Bauknecht, P., Jékely, G. and Hejnol, A. (2017) An ancient FMRFamide-related peptide-receptor pair induces defence behaviour in a brachiopod larva. Open biology, 7.

